# The streptococcal phase-variable Type I Restriction-Modification system SsuCC20p dictates the methylome of *Streptococcus suis* and impacts virulence

**DOI:** 10.1101/2023.03.17.533248

**Authors:** Thomas J. Roodsant, Boas van der Putten, Jaime Brizuela, Jordy P.M. Coolen, Tim J.H. Baltussen, Kim Schipper, Yvonne Pannekoek, Kees C.H. Van der Ark, Constance Schultsz

## Abstract

Phase-variable Type I Restriction Modification (RM) systems are epigenetic regulatory systems that have been identified in numerous human bacterial pathogens. We previously showed that an emerging zoonotic lineage of *Streptococcus suis* acquired a phase-variable Type I RM system named SsuCC20p. The SsuCC20p locus was present in the genome of disease-associated isolates from multiple streptococcal species. This indicates that it is not restricted to *S. suis* and can be acquired through horizontal gene transfer. We demonstrate that SsuCC20p phase-variability relies on a recombinase present within the locus. *In vitro*, only SsuCC20p is responsible for the genome methylation profiles that were detected in the representative zoonotic *S. suis* isolate 861160. In addition, we show that, contrary to previous observations, *hsdS* genes located downstream of the *hsdM* gene and the recombinase gene, can contribute to the SsuCC20p genome methylation profile. SsuCC20p locked mutants expressing a single *hsdS* each showed unique genome methylation profiles. The differential genome methylation of the distinct locked mutants caused phase dependent differences in global gene expression in a growth condition dependent manner. We observed significant differences in virulence between *hsdS* locked mutants in a zebrafish larvae infection model. These data indicate that the streptococcal phase-variable Type I RM system SsuCC20p can impact bacterial virulence via epigenetic regulation of gene expression and potentially contributes to the zoonotic potential of *S. suis*.

**Importance:** Phase-variation contributes to the virulence of bacterial pathogens as it allows a single strain to produce phenotypic diverse subpopulations. Phase-variable Restriction Modification (RM) systems are systems that allow for such phase-variation via epigenetic regulation of gene expression levels. The phase-variable RM system SsuCC20p was found in multiple streptococcal species and was acquired by an emerging zoonotic lineage of *Streptococcus suis*. We show that the phase-variability of SsuCC20p is dependent on a recombinase encoded within the SsuCC20p locus. We characterized the genome methylation profiles of the different phases of SsuCC20p and showed that the differential genome methylation within the phases causes differences in gene expression levels and virulence. Altogether, we show that the acquisition of a phase-variable RM system impacts virulence and can potentially contribute to the zoonotic potential of *S. suis*. Bacterial pathogens can increase their virulence through acquisition of mobile elements containing epigenetic regulatory systems such as RM systems.

## Introduction

*Streptococcus suis* is an opportunistic bacterial pathogen in pigs and an emerging zoonotic pathogen (1). Human infections can lead to meningitis, streptococcal toxic-shock like syndrome and septicemia (2, 3). Human infections are linked to exposure to pigs, such as (occupational) handling of pig (products) or consuming undercooked or raw pig products (2, 4). *S. suis* is classified into serotypes based on capsular polysaccharides (CPS) structure and into sequence types, which in turn are clustered into clonal complexes, based on its genomic background as assessed by multi-locus sequence typing (MLST). Most human infections are caused by *S. suis* serotype 2 of clonal complex (CC) 1, although infections with other serotypes (e.g. serotype 14) and genotypes (e.g. CC20) have also been reported (3, 5).

In the Netherlands, a unique zoonotic serotype 2 CC20 clade has been identified, which is more closely related to the non-zoonotic but virulent serotype 9 CC16 clade than to the zoonotic serotype 2 CC1 clade (5). Three major genomic differences in the accessory genome between CC16 and CC20 have been postulated to contribute to zoonotic potential of CC20 strains (5). These include a capsule switch through acquisition of a serotype 2 CPS locus (i), acquisition of a 89k pathogenicity island, previously identified in Chinese zoonotic outbreak isolates (ii), and acquisition of a 18.5kb prophage region with a complete Type I Restriction Modification (RM) system with phase-variable specificity subunits named SsuCC20p (iii)(5, 6).

Phase-variable RM systems can be found in many pathogenic bacteria and have been shown to regulate bacterial virulence (7). Type I RM systems consist of three host specificity determinants (*hsd*) genes encoding a specificity subunit (S), a modification subunit (M) and a restriction subunit (R) (Fig. 1A) (8). The subunits can form a pentameric complex (HsdS, 2HsdM, 2HsdR) with endonuclease activity and a trimeric complex (HsdS, 2HsdM) with methylase activity (8). HsdS is a DNA-binding protein that determines the target DNA sequences of both complexes. HsdS consists of two separate target-recognition domains (TRD) that each recognize a specific part of the target DNA sequence. The TRDs in the HsdS protein are spatially separated from each other, which in turn separates the TRD-recognized DNA motifs by multiple nucleotides (8). Phase-variable Type I RM systems have multiple TRD regions that can recombine to form different functional HsdS proteins. TRD recombination is mediated via inverted repeats (IRs), which in some systems is (partially) mediated by a recombinase within the same locus (8–10). Unique *hsdS* alleles, when expressed with an *hsdM*, give unique methylation profiles in the genome. The methylation of the genome can affect gene expression by affecting the binding of regulatory proteins, such as transcription factors, to regulatory sequences upstream of genes (11, 12). In this way, the phase-dependent methylation profiles can impact virulence, as was shown for *Streptococcus pneumoniae* (13).

**FIG 1.**
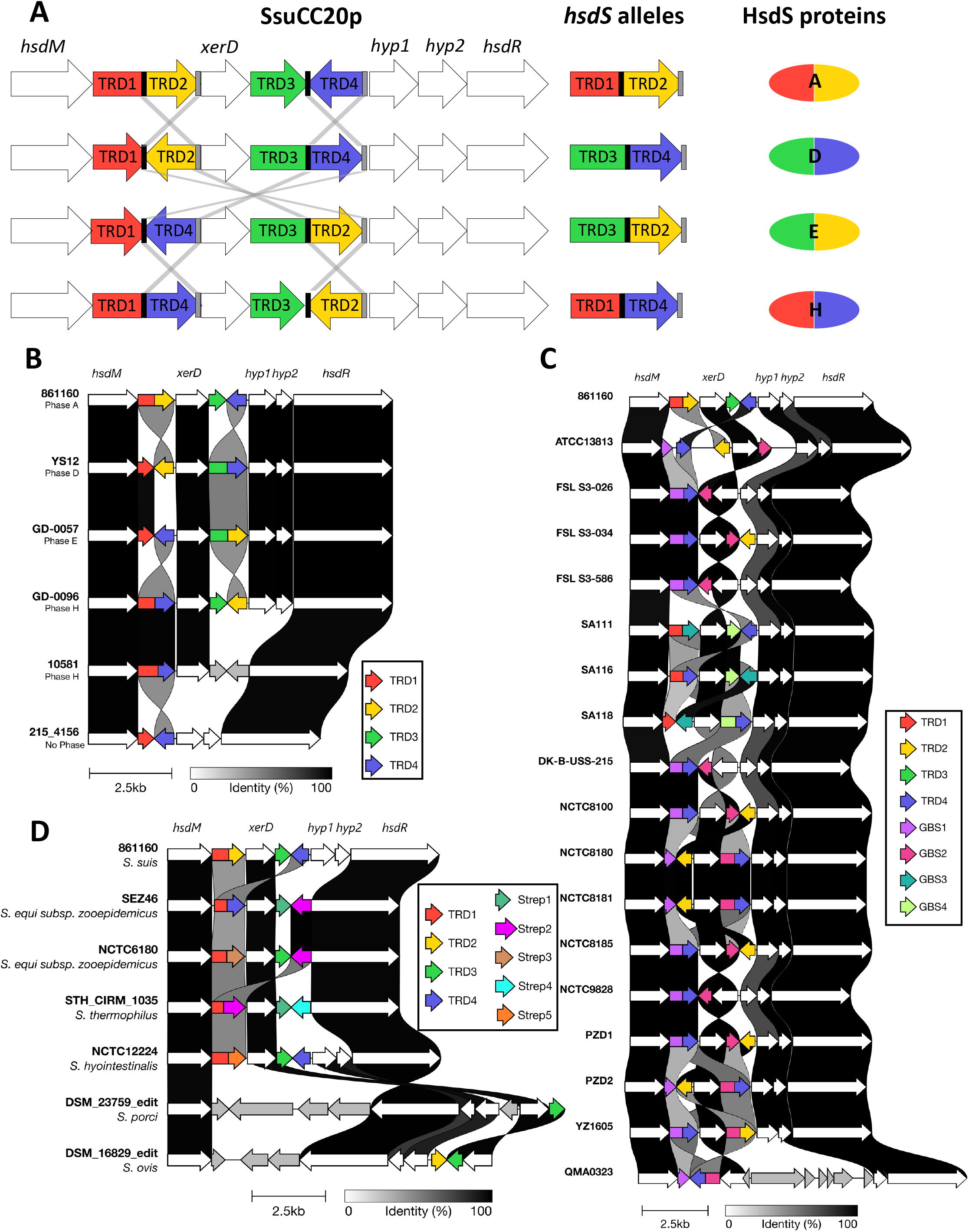
SsuCC20p presence in *S. suis* and other streptococci. (A) Graphical representation of SsuCC20p. Black bars are left inverted repeats and grey bars are right inverted repeats. Grey lines show suggested recombination events. Target recognition domains (TRDs) are numbered 1 to 4. (B) Visualization of unique tblastn hits for SsuCC20p presence in *S. suis*, (C) *S. agalactiae* or (D) other streptococcal species genomes visualized with clinker and clustermap.js, genes with >90% homology are connected by shaded blocks and grey arrows are genes absent in SsuCC20p.

We characterized SsuCC20p and show that SsuCC20p is phase variable, expressed and actively methylates the *S. suis* genome. In a zebrafish larvae infection model of bacterial infection (14–23), isogenic mutants with a single *hsdS* allele and distinct genome methylation profiles differed in virulence.

## Results

### SsuCC20p is present in multiple streptococcal species

The SsuCC20p locus contains not only a *hsdM*, a *hsdR* and 4 TRDs of which two form a functional *hsdS*, but also a site-specific recombinase gene (*xerD*) and two hypothetical proteins (Fig. 1A). Many additional *S. suis* genomes have been sequenced since the discovery of SsuCC20p (5, 6). A curated collection of 1703 *S. suis* genome assemblies (24) supplemented with recently sequenced European zoonotic *S. suis* isolates (J. Brizuela, T. Roodsant, Q. Hasnoea, B. van der Putten, J. Kozakovac, H.C. Slotved, M. van der Linden, I. de Beer, E. Sadowyg, J.A. Saez-Nieto, V. Chalkeri, K. van der Ark, C. Schultsz, submitted for publication) was searched for the presence of the SsuCC20p locus (Fig. 1A; supplemental material Table S1). A total of 22 isolates carried the complete SsuCC20p locus. Two isolates (215_4156 and 216-4157) missed *xerD*, TRD2 and TRD3 and had lower homology (78% and 79%) with hypothetical protein 2. One isolate (10581) missed both hypothetical proteins, TRD2 and TRD3 (Fig. 1B, supplemental material Table S2). Additional strains in which SsuCC20p was identified include two human isolates from Germany (1) and the Czech Republic (1), and four pig isolates from Australia (2), the US (1) and Denmark (1). Besides CC20 isolates, SsuCC20p was found in one strain belonging to CC25 (1) and in three strains not yet assigned to a CC. Four different *hsdS* alleles were found in the assemblies of *S. suis* genomes, which encoded four unique HsdS proteins (Fig. 1A,B). Allele H was found most frequently within CC20 genomes (n=10), followed by A (n=6) and E (n=5). Allele D was only found within one Chinese ST17 isolate. All except one isolate were associated with disease (supplemental material Table S2).

SsuCC20p is located on a prophage region and is therefore likely acquired via horizontal gene transfer (6). To identify other species that carry the same Type I RM system, we searched the NCBI Refseq Genomes Bacterial Database for the presence of each individual gene within the SsuCC20p locus. SsuCC20p was restricted to streptococci and most hits (17/23) were found in *Streptococcus agalactiae*. The *hsdS* alleles found in *S. agalactiae* lacked multiple TRDs compared to *S. suis hsdS* alleles A, D, E or H (Fig. 1C; supplemental material Table S3). All but one *S. agalactiae* strain had a functional *hsdS* comprised of two TRDs and five strains had three instead of four TRDs within the locus. In these same five *S. agalactiae* strains, the orientation of *xerD* was reverted compared to the rest of the locus and these strains had lost the IRs identified in *S. suis* SsuCC20p. SsuCC20p was also identified in five other streptococcal species, but with more genomic differences than found for the *S. agalactiae* isolates, including loss of genes, acquisition of a new TRD or large intergenic insertions (Fig. 1D; supplemental material Table S3; supplemental material Table S3). In case the host health state was reported for the isolate carrying this type I RM system (10/23), the isolate was associated with disease (supplemental material Table S3).

### SsuCC20p is phase-variable

The presence of a site-specific recombinase (*xerD*) and TRDs flanked by IRs within a single locus, combined with the presence of multiple *hsdS* alleles in different genome assemblies suggest that SsuCC20p is phase-variable. Whilst four different alleles of SsuCC20p have been identified in CC20 *S. suis* genomes, the presence of these different alleles within a single isolate has not been demonstrated yet. We chose the zoonotic ST20 isolate 861160 as our model strain because a single contig assembly of its genome is available (25). We identified and quantified the allelic variants after overnight growth in THY in a FAM-labelled PCR with a subsequent endonuclease digestion and fragment analysis (Fig. 2A), as previously used to quantify phase-variable *hsdS* alleles (9, 13, 26). The most prevalent allelic variant was *hsdS* A, in accordance with the *hsdS* allele present in the 861160 genome assembly (GCA_902702745), followed by *hsdS* E and *hsdS* H (Fig. 2B,C). Allele D was undetected, in line with its absence in the assembled genomes of ST20 isolates. Using long read sequencing, we corroborated this allele distribution within 861160. In total, 16 nanopore reads and 205 single-molecule real time (SMRT) sequencing reads spanned the complete *hsdS* locus and both methods showed a comparable allele distribution (Fig. 2D,E).

**FIG 2.**
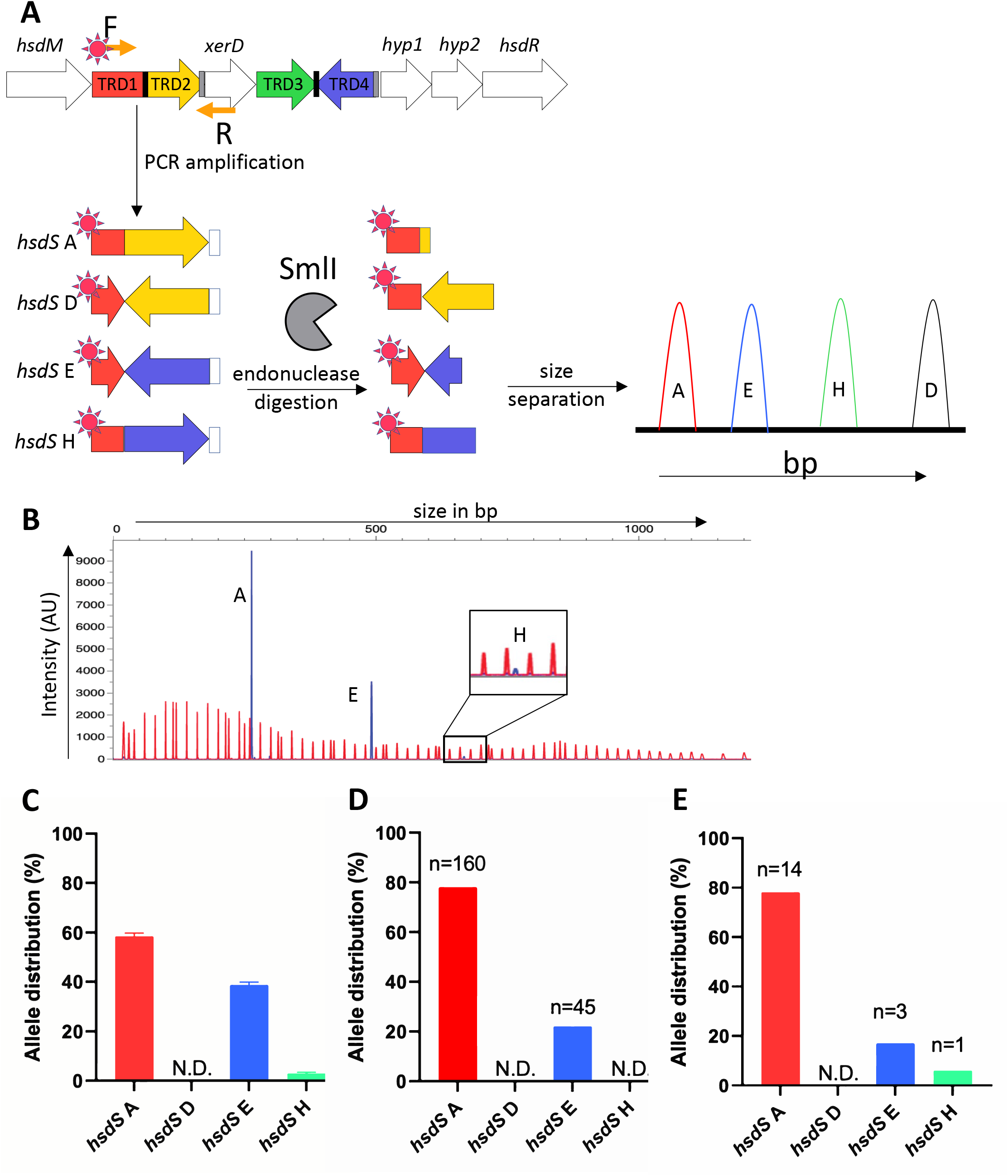
SsuCC20p is phase-variable and three alleles can be found within a single isolate. (A) Graphical representation of the *hsdS* allele quantification by FAM-labelled PCR product endonuclease digestion and fragment analysis. (B) Representative example of WT 861160 *hsdS* allele identification, red peaks are the LIZ1200 size marker and blue peaks the FAM-labelled fragments. (C) *hsdS* allele distribution was computed by measuring the relative area under the curve of the different *hsdS* alleles using PeakScanner v3.0 in three biological replicates. (D) *hsdS* alleles were quantified using PacBio or (E) Oxford Nanopore long read sequencing data, read count per allele is indicated above each bar. N.D. is Not Detected, AU is Arbitrary Units.

### SsuCC20p is expressed and differentially methylates the genome

In previously identified Type I RM systems, the *hsdS* allele that is not located directly downstream of the *hsdM* gene is silent, because it is encoded on the opposite strand, lacks a start codon, or lacks a promoter (27). In SsuCC20p, a complete and functional *hsdS* is predicted not only directly downstream of the *hsdM* gene in allele A and H, but also downstream of *xerD* in allele D and E (Fig. 1A). To investigate if both TRD1 (allele A and H) and TRD3 (allele D and E), as well as the other genes of SsuCC20p are transcribed in 861160, we performed RT-PCR. All genes of SsuCC20p were expressed in 861160 when grown into logarithmic phase in THY (Fig. 3A). Gene expression quantification by RT-qPCR showed that the expression of TRD1 and TRD3 was similar, but both TRDs had lower expression levels than *hsdM, hsdR* and *xerD* (Fig. 3B).

**FIG 3.**
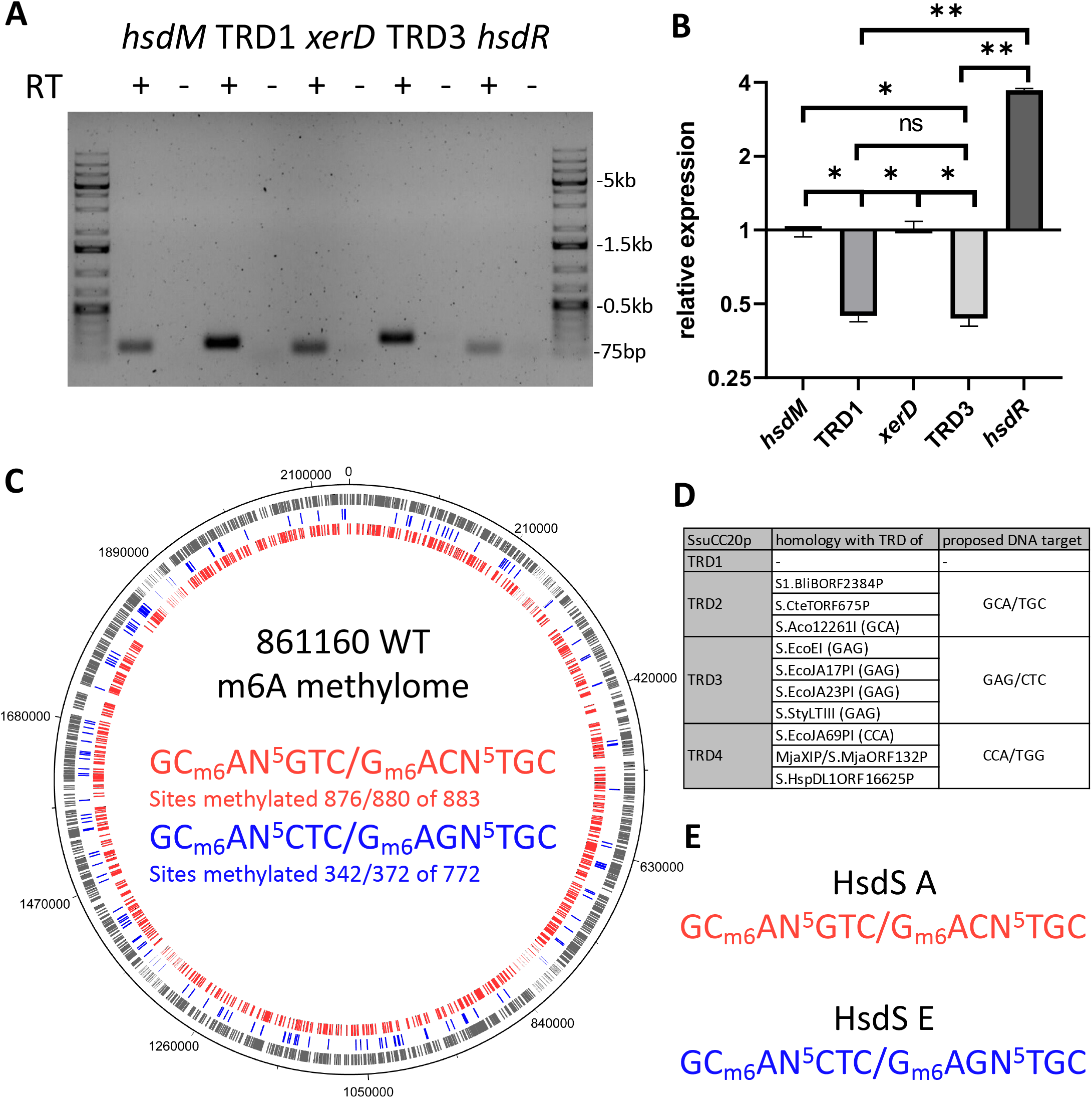
*hsdS* alleles downstream and upstream of *xerD* are expressed and methylate the genome. (A) Expression of SsuCC20p genes was verified by PCR on reverse transcribed RNA, using gene specific qPCR primers. PCR products were run on a 1% agarose gel. RT + or – indicate the presence of reverse transcriptase in the reverse transcription reaction, marker lanes were loaded with the 1kb+ ladder. (B) Expression quantification of genes present in SsuCC20p. Genes were normalized against *proS* and *gdH* (57,58), expression levels are shown relative to *hsdM*. Error bars represent means ± standard deviations from three biological replicates. Asterisks denote significance difference by one-way ANOVA with Tukey’s multiple comparisons (α<0.05), * p<0.001 and ** p<0.0001. (C) 861160 WT methylome, m6A methylated sites in genome are indicated on the genome with the colour corresponding to the DNA motif, grey indicates sites that have either of the two DNA motif sites. (D) Homology of SsuCC20p TRDs to other Type-I RM system *hsdS* genes, identified DNA target sequences are indicated between brackets, for details see supplemental material Fig. S2. (E) proposed target DNA motif of HsdS A and HsdS E.

To determine if SsuCC20p methylates the genome, the methylome of wildtype 861160 grown in THY was compared with a *ΔhsdS* strain lacking each of the 4 TRDs and *xerD* using PacBio HiFi sequencing. In the wildtype, two m6A methylation profiles were identified (GC_m6_AN^5^GTC/G_m6_ACN^5^TGC and GC _m6_AN^5^CTC/G _m6_AGN^5^TGC) with the Type I RM system characteristic bipartite build-up of the target DNA motifs (8), and both methylation profiles were absent in the Δ*hsdS* strain (Fig. 3C). Almost all potential sites of GCAN^5^GTC/GACN^5^TGC (99%/100%) within the full genome were methylated, but the GCAN^5^CTC/GAGN^5^TGC sites were only partially methylated (44%/48%). The partial methylation of the GCAN^5^CTC/GAGN^5^TGC sites was also observed in a second PacBio HiFi sequencing run on newly isolated genomic DNA, in which 37%/30% of the sites were methylated (supplemental material Fig. S1). The methylated sites had overlap between both runs, but for each run also unique m6A methylations sites were detected (supplemental material Fig. S1).

To assign the *hsdS* alleles found in WT 861160 (A, E and H) to the two identified methylation profiles, we searched the genome for homologues of the *hsdS* alleles for which DNA targets were identified, using InterPro (v5.55-88.0, default settings)(28), which gave hits for TRD2, TRD3 and TRD4 (supplemental material Fig. S2). Based on the homology we predict that TRD2, TRD3 and TRD4 recognize GCA/TGC, GAG/CTC and CCA/TGG respectively (Fig. 3D). In 861160 WT, *hsdS A* and *hsdS* E are the dominant alleles and responsible for the observed methylation profiles. Both *hsdS* alleles have TRD2 which likely recognizes GCA/TGC (Fig. 3D), but *hsdS* E also has TRD3, which based on homology, likely recognizes GAG/CTC (Fig. 3D). Combining the homology data with the observed methylation profiles and dominant *hsdS* alleles, we propose that GCm6AN^5^GTC/G _m6_ACN^5^TGC is methylated by HsdS A and GC_m_6AN^5^CTC/G_m_6AGN^5^TGC by HsdS E (Fig. 3E).

### Phase-variability of SsuCC20p is *xerD* dependent

In *S. pneumoniae*, the site-specific recombinase (*creX*) present in the Type I RM system SpnIII is essential for the TRD recombination between the small IRs (15bp), but not for TRD recombination between the larger IRs (85bp and 333bp) (9). The IRs within SsuCC20p are 14bp, therefore we expected that TRD recombination in SsuCC20p is mediated by *xerD* present in the locus, which was tested in Δ*xerD* mutants in 861160. After three sequential subcultures, to allow for recombination, the *hsdS* alleles present in the Δ*xerD* mutants and WT strain were assessed using the previously described endonuclease digested FAM-labelled PCR product fragment analysis. In the WT, all three *hsdS* alleles were detected (Fig. 4A), but in the Δ*xerD* mutants only a single *hsdS* allele per mutant was identified, which was *hsdS* A, E or H depending on the *hsdS* allele present in the genome upon mutating *xerD* (Fig. 4B-D). Long read sequencing confirmed the fixed state of the *hsdS* alelles in the Δ*xerD* mutants. In addition, SMRT and HiFi sequencing showed that the kanamycin cassette used to create the mutants, disrupted SsuCC20p genome methylation (supplemental material Table S4).

**FIG 4.**
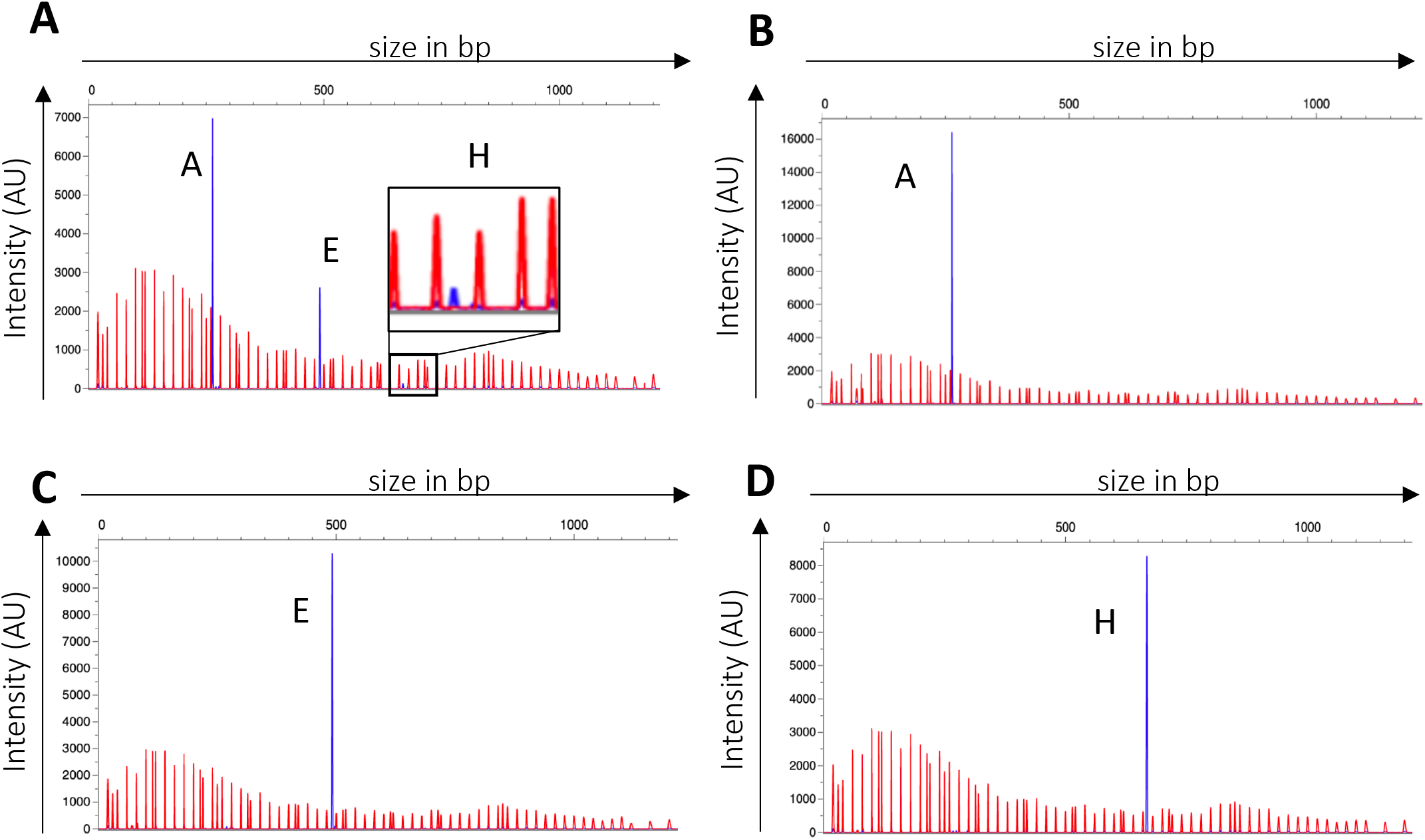
SsuCC20p phase-variability is *xerD* dependent. (A) Representative example of WT (A) and Δ*xerD* (B-D) *hsdS* allele identification after subculturing in THY broth. Red peaks are the LIZ1200 size marker and blue peaks the FAM-labelled fragments, *hsdS* allele identification was analyzed with PeakScanner v3.0 in three biological replicates. Δ*xerD* mutants showed a single *hsdS* allele, which was *hsdS* A, E or H depending on the *hsdS* allele present in the genome upon mutating *xerD*.

### *hsdS* alleles give unique methylomes

To test the predicted *hsdS* allele assignment to the observed methylation profiles in the WT 861160 strain, locked mutants (LM) were constructed in which the silent TRDs and *xerD* were replaced by an erythromycin resistance cassette so that the LMs express only a single *hsdS* allele (Fig. 5A). In all LMs, the expression of *hsdM, hsdS*, and *hsdR* was confirmed using RT-PCR (supplemental material Fig. S3). SMRT and HiFi sequencing of the mutants showed that each LM expressed a single unique m6A methylation profile (Fig. 5B) and that on homology based predicted methylation profiles for *hsdS* A and E were correct (Fig. 3C and 5B). Additionally, we identified the methylation profile of *hsdS* H (CC _m6_AN^5^GTC/G _m6_ACN^5^TGG). In contrast to the *hsdS* E methylation profile in WT, all potential methylation sites for *hsdS* E were methylated. Of note, genome analysis showed that the DNA sequences of the LMs were identical, excluding their unique *hsdS* alleles, but all had acquired a SNP in the *cps2F* gene (371A>G) compared to their parent strain.

**FIG 5.**
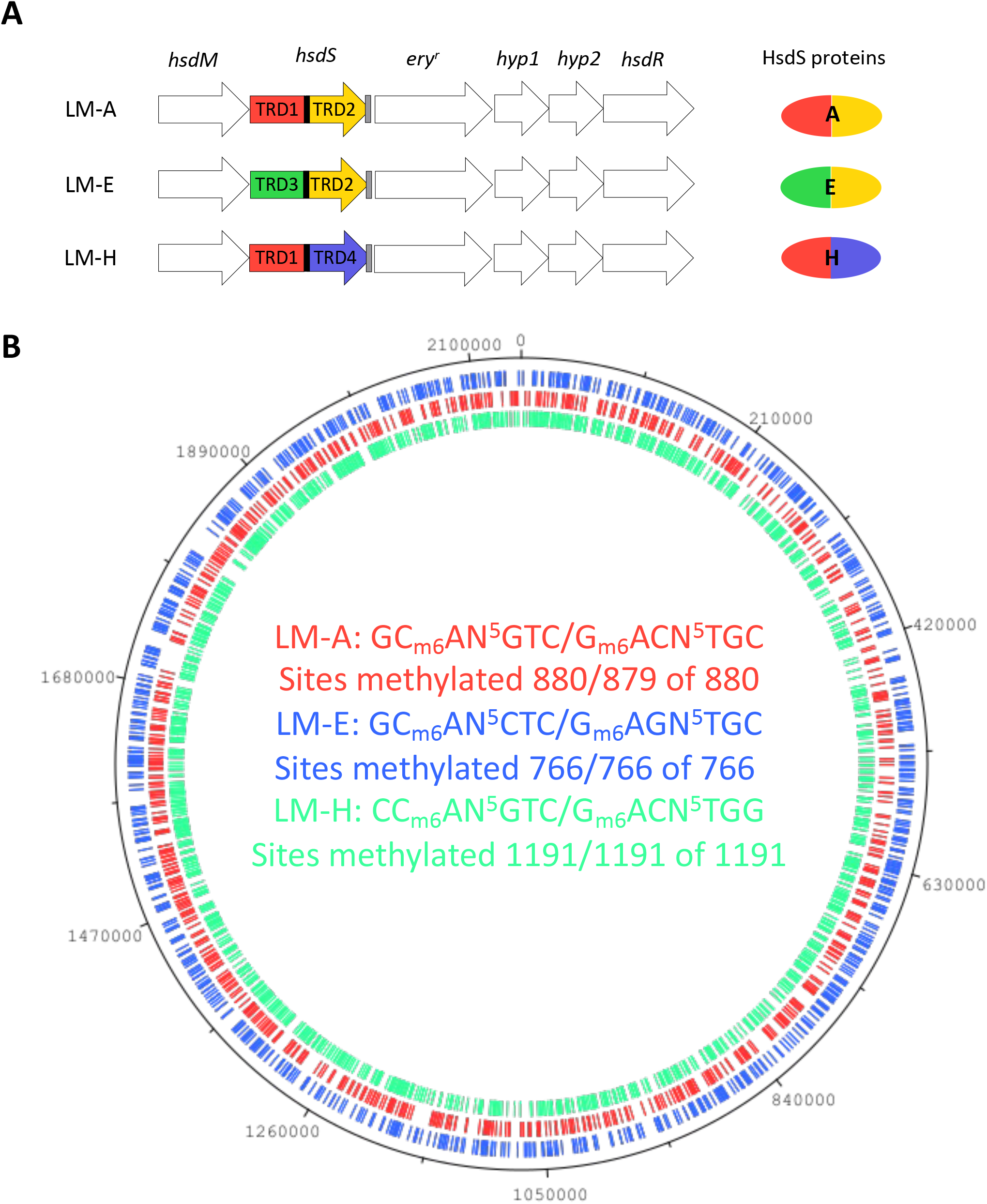
Locked mutants (LMs) have unique m6A methylation patterns. (A) Graphical representation of the SsuCC20p locus in the LMs. (B) The LM methylation patterns were plotted on the 861160 WT genome, m6A methylated sites in the genome are indicated on the genome with the colour corresponding to the DNA motif.

### LMs differ in gene expression and virulence in a zebrafish larvae infection model

The unique genome methylation profiles of the LMs can result in differentially expressed genes, as shown for CPS biosynthesis genes in *S. pneumoniae* (13). Thus, we aimed to identify genes that are differentially expressed in the LMs and screened the 861160 genome sequence for promoter sequences which contain either of the three identified methylation profiles. Using multiple promoter predictor tools, we identified 24 genes (LM-A n=7, LM-E n=6, LM-H n=11) carrying a methylation profile in their promotor sequence region. The promoter sequence regions of the identified genes did not show homology. Next, differential expression was assessed by RT-qPCR for logarithmic cultures grown in THY or human serum for the identified genes and two CPS biosynthesis genes. None of the genes differed in expression between the LM when bacteria were grown in THY (Fig. 6A). When grown in human serum, 8 genes were significantly differential expressed in LM-H compared to LM-A and LM-E, of which three had at least a 2-fold difference (Fig. 6B). The serine/threonine transporter (*sstT*, RS03300) was 1.99-2.06 fold higher expressed in LM-H than LM-A or LM-E. The expression of the pseudo gene RS04125 was 5.40-6.28 fold higher in LM-H than LM-A or LM-E. LM-H showed a 2.54-2.97 fold higher expression of the DUF5590 domain-containing protein RS04145, which is at the start of an operon that also encodes as a pyridoxal phosphate-dependent aminotransferase (RS04150) and an asparagine--tRNA ligase (RS04155).

**FIG 6.**
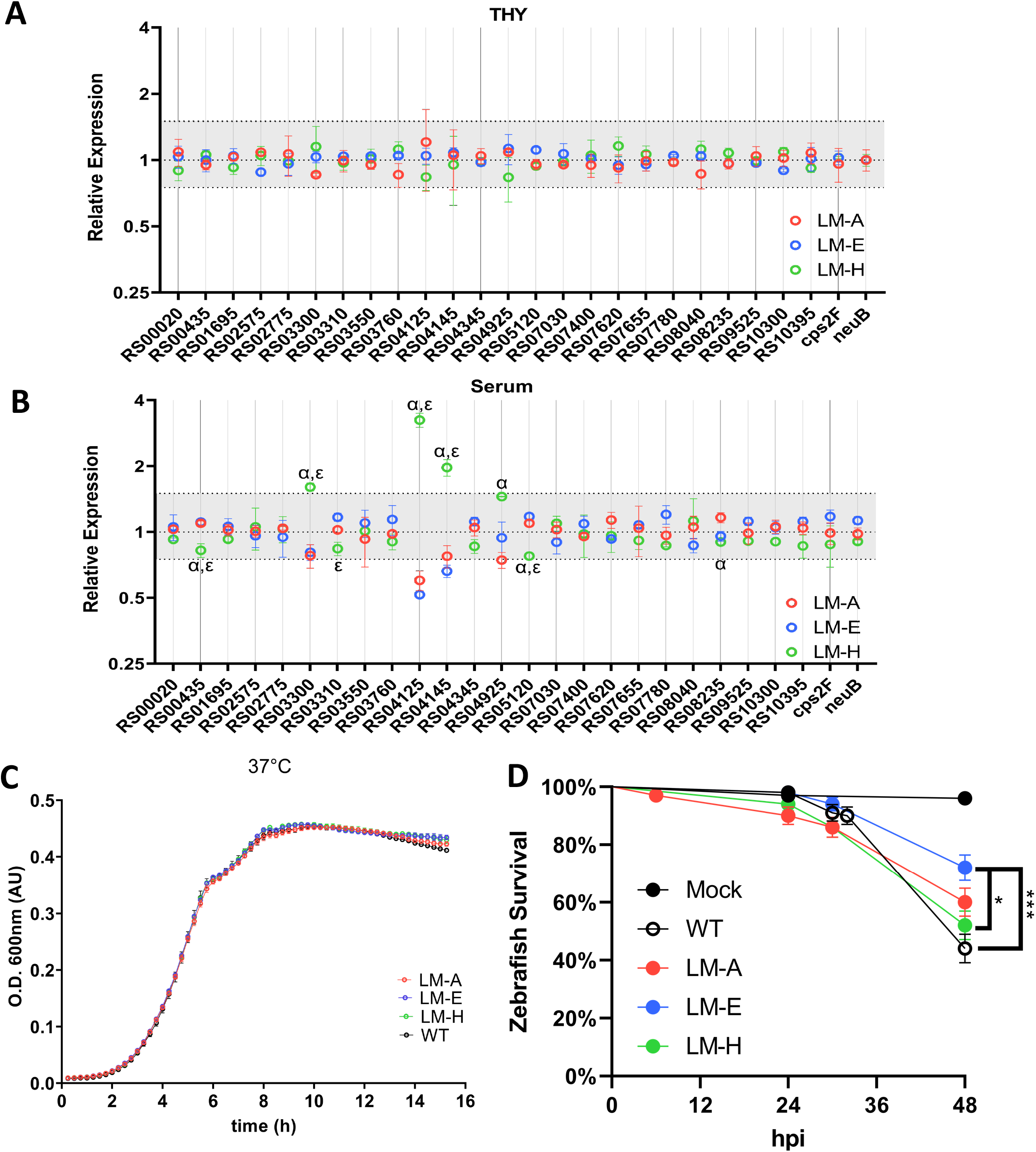
Locked mutants of SsuCC2Op show differential gene expression and different virulence in a zebrafish larvae infection model. (A,B) Gene expression quantification by RT-qPCR in LM-A,LM-E and LM-H grown in THY (A) or human serum (B) (n=3). Genes are indicated by locus tags according to the published 861160 genome assembly and annotation (GCF_902702745.1, (24)). Gene expression was normalized using *gdH* and *proS*, and expression per gene was plotted relative to the geometric mean expression of that gene by all LMs combined (means±SD). Relative expression between 0.75 and 1.5 was shaded grey. Statistics were performed on normalized data using a one-way ANOVA with Bonferroni correction for multiple testing, significant differences between LM-H (p<0.05) compared to LM-A and LM-E are indicated with α and ε respectively. (C) Growth of locked mutants in THY at 37°C was measured by optical density (O.D.) at 600nm in arbitrary units (AU) for 16 hours (h). (D) Zebrafish Larvae (72 hours post fertilization) were infected via yolk sac injection with 2700 CFU of *S. suis* and survival was followed for 48 hours post infection (hpi). Per group, 20 larvae were infected in 5 separate experiments. Data from individual experiments were pooled and statistical difference was determined using a log-rank (Mantel-cox) test with a Bonferroni correction for multiple testing, * p<0.05, ** p<0.01 and ***p<0.001.

The unique genome methylation profiles of the LMs could affect *S. suis* antimicrobial resistance, growth or virulence (13). The LMs did not differ in growth rate in THY at 37 °C or in antimicrobial susceptibility to several antibiotics (Fig. 6B, supplemental material Fig. S4A). The potential difference in virulence of LMs was assessed in a zebrafish larvae infection model, which has been used to study the virulence of *S. suis* and other *Streptococcus* species before (14–16, 19, 29). In the zebrafish larvae infection model, LM-E was less virulent causing a higher survival of zebrafish than LM-H or WT, LM-A did not significantly differ from LM-E, LM-H or WT (Fig. 6C). This difference could not be attributed to difference in growth rate at the zebrafish larvae incubation temperature (28 °C) (supplemental material Fig. S4B).

## Discussion

Epigenetic regulation of bacterial gene expression by phase-variable RM systems plays an important role in host-microbe interactions, including bacterial pathogenesis (7, 13, 30–33). Here, we characterized the phase-variable Type I RM system SsuCC20p that is encoded on a mobile element and is found in multiple streptococcal species including a zoonotic *S. suis* lineage. The SsuCC20p *hsdS* allele dependent genome methylation impacted *S. suis* gene expression and virulence in a zebrafish larvae infection model for bacterial pathogenesis.

In contrast to previously characterized phase-variable Type I RM systems, we demonstrate that additional *hsdS* alleles further downstream of *hsdM* are not always silent. To the best of our knowledge, in all phase-variable Type I RM systems described so far, only the *hsdS* allele directly downstream of the *hsdM* is transcribed and involved in genome methylation (10, 13, 26, 27, 30, 31, 34). Therefore, it was suggested that the other TRDs or *hsdS* within the locus are merely present to function as DNA templates for recombination (34). However, *hsdS* E of SsuCC20p is transcribed and methylated the genome despite not being located directly downstream of *hsdM*.

Most CC20 isolates including 861160 encode three Type I RM systems (SsuCC20p, SsuPORF1588P and SsuPORF1273P) (6), but only SsuCC20p actively methylates the genome. In many prokaryotes, multiple (Type I) RM systems methylate the genome simultaneously (10, 34, 35). Closer inspection of SsuPORF1588P showed that both *hsdS* have only a single TRD instead of the two TRDs that almost all functional Type I RM *hsdS* encode (supplemental material Fig. S5A)(8), thus we speculate that SsuPORF1588P is not functional in methylation. SsuPORF1273P is phase-variable (26) and the methylation profile of its two dominant alleles (A and B) has been solved by SMRT sequencing of *E. coli* overexpressing the *S. suis* methyltransferases, giving CC_m6_AN^8^CTT for allele A and CCm6AN^6^DNH (D=A/G/T, H=A/C/T and N=A/C/G/T) for allele B (27). SsuPORF1273P expression when grown overnight on BHI plate has been demonstrated, but genome methylation in *S. suis* has not been shown yet (26). In 861160, SsuPORF1273P appears functional and carries *hsdS* allele C (supplemental material Fig. S5B), thus SsuPORF1273P should be able to methylate the genome when expressed. The different culture condition (THY broth vs BHI plates (26)) could explain the absence of SsuPORF1273P methylation in our hands, although other mechanisms regulating transcription or translation of SsuPORF1273P could also be involved. The genome methylation by only a single Type I RM system in the presence of multiple encoding RM systems, due to differences in transcription levels under specific culture conditions, has been observed in *Streptococcus pyogenes* as well (36).

Our query for SsuCC20p in *S. suis* isolates identified 4 *hsdS* alleles, but only 3 could be detected within the 861160 isolate under the used experimental conditions. Strain YS12 (ST17), that has the *hsdS* D allele in its genome assembly, belongs to a different ST than the strain used in our experiments (ST20). Based on the 861160 genome, the undetected *hsdS* D allele could be formed via *xerD* mediated recombination. In *S. pneumoniae*, the absence of specific *hsdS* alleles within an isolate under specific culture conditions was observed (13). Thus, we speculate that our experimental conditions do not sustain a ST20 strain with the *hsdS* D allele, or that this strain is outcompeted by strains expressing the other three alleles bringing its abundance below detection limit.

The partial *hsdS* E and absent *hsdS* H methylation detected in 861160 WT are likely caused by the consensus approach used to extract the methylation profile from the SMRT sequencing reads, explaining why all potential methylation sites are methylated in LM-E and LM-H. A similar observation was made for the Spn556I in *S. pneumoniae* ST556 WT, in which all potential sites were methylated for the dominant *hsdS* allele but only a fraction, or none, of the potential sites were methylated for the minor *hsdS* alleles (10). Methylated nucleotides affect the interpulse duration measured during SMRT sequencing, which is used to identify methylation profiles (11). We used a consensus approach for methylation detection, in which SMRT reads are aligned to obtain the mean interpulse duration (IPD). We speculate that the IPD value of *hsdS* E or H methylated reads are averaged with the dominant *hsdS* A methylated reads to a level that the potential methylation sites are classified as unmethylated, leading to the partial *hsdS* E and absent *hsdS* H methylation profile detected in 861160 WT.

Different SsuCC20p *hsdS* alleles result in unique genome methylation profiles and impact both the gene expression during *in vitro* culture and the virulence in a zebrafish larvae infection model of a zoonotic CC20 isolate. Despite a similar innate immune system and its proven usefulness to study human pathogens (16–18, 20–23), the zebrafish model has limitations. For example, the absence of an adaptive immune system, anatomical differences and a lower body temperature, thus findings should carefully be extrapolated to human infections (23, 37, 38).

The impact of genome methylation on the transcriptome is dependent on culture conditions. When grown in serum, 8 genes were differentially expressed in the LMs, while they showed similar expression when the LMs were grown in THY. A similar observation was made in *Bacteroides fragilis*, differentially expressed genes or the fold change of genes differentially expressed within LMs differed between *in vivo* and *in vitro* culture conditions (31). For the three most differentially expressed genes in *S. suis*, the only characterized gene is *sstT*. *sstT* is responsible for the uptake of serine and threonine (39), which could increase bacterial fitness. In *S. pyogenes, sstT* was essential for bacterial survival during skeletal muscle necrotizing myositis (40). The differential expression of other genes of the LMs can likely explain the observed difference in virulence in the zebrafish model. A less targeted approach such as RNA sequencing is more likely to identify all genes differentially expressed in the LMs, including genes that are indirectly affected by genome methylation (as reviewed in (11)).

The importance of epigenetic regulation by phase-variable RM systems has been demonstrated in many bacterial pathogens (7). Multiple lineages of *S. suis* carry phase-variable RM systems of which some are restricted to specific lineages (26). We characterized a phase-variable Type I RM system in the newly identified zoonotic CC20 lineage. Interestingly, pathogenicity in bacteria, including *S. suis* and the CC20 lineage, has been correlated with a reduction in gene content (5, 41, 42), including the loss of transcriptional regulators in virulent *S. suis* isolates (42). We propose that phase-variable RM systems offer transcriptional flexibility via epigenetic regulation, allowing a reduction of other transcriptional regulators in the genome while maintaining phenotypic diversity within the isolate that is needed to thrive or survive in new environments. Thus, explaining the co-occurrence of reduced gene content and phase-variable RM system acquisition in virulent and zoonotic *S. suis* lineages. Here, we characterized the phase-variable Type I RM system SsuCC20p and demonstrated its impact on *S. suis* virulence via differential genome methylation. Moreover, we find that SsuCC20p is predominantly found in disease associated isolates in *S. suis* and other streptococcal species, arguing for a potential role of SsuCC20p in *S. suis* virulence.

## Material and Methods

### SsuCC20p identification in *S. suis*

A custom made ABRicate v1.0.1 (https://github.com/tseemann/abricate) database containing the nucleotide sequences of the five genes and four TRDs present within the SsuCC20p locus (Supplementary material Table S1) was run (blastn) against a curated collection of 1703 *S. suis* genome assemblies supplemented with 46 genome assemblies of recently sequenced European zoonotic *S. suis* isolates (24) (J. Brizuela, T. Roodsant, Q. Hasnoea, B. van der Putten, J. Kozakovac, H.C. Slotved, M. van der Linden, I. de Beer, E. Sadowyg, J.A. Saez-Nieto, V. Chalkeri, K. van der Ark, C. Schultsz, submitted for publication) using default settings. For the identification of SsuCC20p in other bacterial species, the ABRicate database was translated to protein and used to search with tblastn against the NCBI Refseq Genomes Bacterial Database, excluding *S. suis* using default settings. Hits within the same contig with a minimal identity of 90% for the non-phase-variable genes *hsdM, xerD* and *hsdR* were selected for genomic region extraction. The extracted genomic regions containing SsuCC20p were aligned using clinker and clustermap.js (43).

### Bacterial strains and culture conditions

The *S. suis* strains used are listed in the supplemental material (Table S4). *S. suis* was grown in Todd-Hewitt Broth supplemented with 0.5% yeast extract (THY) at 37 °C, supplemented with 200 μg/ml kanamycin or 2 μg/ml erythromycin when required.

### *hsdS* allele quantification

SsuCC20p *hsdS* allele quantification was adapted from previously published protocols for other phase-variable RM systems (13, 26). In short, FAM labelled PCR fragments were amplified from genomic DNA using the primers ‘5-6FAM-CTGGAGGGTGTTCTAATGATG-3’ and ‘5-CCGCTCGCTATTTCCTA-3’. Fragments were gel purified and enzymatically digested with SmlI (NEB) for 1h at 55 °C. Digested fragments were diluted in HiDi formamide and ran on the ABI 3730XL DNA analyzer using the LIZ1200 size standard. *hsdS* alleles were quantified by measuring the area under the curve using Peak Scanner (v3.0).

### Cloning

Mutants listed in the supplemental material (Table S4) were created by homologous recombination using peptide induced competence and mutator fragments containing an antimicrobial resistance marker (kan^r^ (44) or ery^r^ (45)) with >500 bp homologues flanking regions as previously published (46). Briefly, 5 μM of the ComS13-21 competence inducing peptide and 1-5 μg of mutator fragment was added to *S. suis* grown to an optical density (OD, 600 nm) of 0.05 and incubated for 2 h at 37 °C. Hereafter, transformants were selected on THY agar plates containing corresponding antibiotics. Mutator fragments were generated using overlapping PCR, or by PCR amplification of a synthesized gblock (IDT DNA technologies) for the Δ*hsdS* mutant, with the primers listed in the supplemental material (Table S5). Mutants were confirmed by whole genome sequencing.

### SMRT sequencing and methylome analysis

An overnight culture of *S. suis* was diluted 500x in fresh pre-warmed THY and incubated at 37 °C until it reached an OD of 0.30-0.45. Genomic DNA was isolated using the Wizard genomic DNA purification kit (Promega) according to manufacturer’s protocol. DNA was quantified with Qubit and DNA integrity was assessed on a 0.7% agarose gel. For detailed sequencing methods see supplemental materials. Briefly, sequencing libraries were constructed from sheared genomic DNA ( 5-20kb) and sequenced on a SMRT Cell 8M using a PacBio sequel II(e) system (Pacific Biosciences).

### Oxford Nanopore Technologies sequencing

Genomic DNA of a *S. suis* culture grown overnight in THY was isolated using the MagAttract high-molecular-weight DNA extraction kit (Qiagen). DNA was quantified with Qubit and DNA integrity was assessed on a 0.7% agarose gel. Library preparations and Nanopore sequencing was performed as previously described (25).

### Potentially LM differentially expressed gene identification

The 861160 genome was manually screened for genes that had either of the three methylation profiles within 100 bp upstream of their start codon in Artemis (25, 47), and the 81 bp directly upstream were extracted. Promoter sequences were identified within these 81 bp with iPro70-FMWin (48), BRPOM (49), iPromoter-2L(50), promotech (51) and previously identified *S. suis* promoters (52), using default settings. Genes downstream of predicted promoter sequences (based on BPROM or Promotech) that had overlap with either of the three methylation profiles were selected for RT-qPCR.

### Serum collection

Blood was collected from healthy volunteer donors after informed consent into CAT serum clot activator tubes (Greiner Bio-One). After 30 min, serum was separated from clotted blood by centrifugation for 15min at 2200 xg. Serum of 4 donors was pooled and stored at −70 °C until used.

### RNA isolation and RT-(q)PCR

For growth in serum the protocol was adapted from (53), an overnight culture was diluted to an OD of 0.4 and diluted 10x in serum, then grown for 150 min at 37 °C at 5% CO_2_. For growth in THY, an overnight culture of *S. suis* was diluted 500x in fresh pre-warmed THY and incubated at 37 °C until it reached OD of 0.30-0.45. Culture (10 mL) was pelleted (10 min, 4000 xg, 4°C) and resuspended in 1.0 mL Trizol (Invitrogen). Cells were lysed by beat-beating in a MagnaLyser (30 s, 6000 speed) using 0.1 mL zirconium sand and 3×2mm glass beads, with one round of beat-beating for THY cultures and five round for serum cultures. RNA was purified from lysed cells using the direct-zol RNA miniprep kit (Zymo Research). Genomic DNA was removed with the TURBO DNA free kit (Invitrogen) in presence of SUPERase RNAse inhibitor (Invitrogen). cDNA was made from RNA (2.0 μg for THY and 0.6 μg for serum) with Superscript IV first strand synthesis kit (Invitrogen) using random hexamers. Gene expression was assessed by qPCR using primers listed in the supplemental material Table S6 and the GO Taq PCR kit (Promega). Gene expression was quantified using SYBR Green (Roche) in a CFX96 or CFX384 (BioRad) and analyzed using LinRegPCR (54, 55), using *proS* and *gdH* as reference genes (56, 57). All kits were used according to manufacturer’s protocol.

### Growth Curve

Overnight cultures were diluted to an OD of 0.02 in fresh THY broth and 150 μL was distributed in a 96 well flat bottom plate. Bacterial growth at 37 °C or 28 °C without shaking was monitored by measuring the OD every 10 min for 16h in Biotek Synergy H1 plate reader. The assay was performed *in triplo* with three biological replicates.

### Antimicrobial susceptibility testing

E-tests (Oxoid) and disc diffusion assays (BD) were performed as described by the EUCAST guidelines. In brief, strains were grown overnight on blood agar plates, and their initial concentration adjusted using MacFarland standards. Bacterial lawns were incubated with the appropriate E-test stripes or antimicrobial discs. Disc zone diameter and MIC were compared with wildtype and LM.

### Zebrafish larvae infection

Zebrafish larvae infections were performed as previously published (15) with minor adjustments. *S. suis* glycerol stocks were prepared to ensure consistent inocula in between experiments. *S. suis* cultures grown to an OD of 0.4 were frozen in 15% glycerol and stored at −80 °C. For each experiment, stocks were thawed and washed once with PBS, before resuspension in injection buffer (0.125% phenol red; Sigma P0290 in PBS) to obtain a bacterial suspension of 2700±300 CFU/nL. Bacterial inoculum was quantified each infection experiment by serial dilution and plating on blood agar plates.

Adult zebrafish were handled in compliance with the local animal welfare regulations approved by the local animal welfare committee (DEC) and were maintained according to standard protocols (www.zfin.org). Experiments with zebrafish embryos younger than 5 days post fertilization did not need ethical permission. Eggs were harvested within 1 hour after spawning and kept in E3 medium (www.zfin.org) at 28 °C before further use. At 72 hours post fertilization (hpf), vital zebrafish larvae were anesthetized with 0.4% Tricaine in H_2_O and injected in the yolk sac with 1 nL of inoculum or injection buffer (mock). Injected larvae were kept grouped (20 per petri dish) in E3 medium at 28 °C. Zebrafish survival was assessed by visual assessment of a heartbeat at 6, 24, 30 and 48 hours post infection, and diseased zebrafish larvae were removed.

### Statistical analysis

All statical analyses were performed using GraphPad Prism (v9.3.1) and statistical test and p values are indicated in figure legends.

## Data availability

Raw sequencing data and genome assemblies have been deposited in ENA under the accession numbers listed in the supplemental material Table S4.

## Ethical approval

Blood was collected for this study according to the guidelines of the Declaration of Helsinki and approved by the Institutional Review Board (or Ethics Committee) of AMC-UVA (BACON 1.7 12-07-2017).

## Acknowledgements

We thank Ted Bradley in the Core Facility Genomics (Amsterdam UMC) for conducting the DNA fragment analysis, Irma Schouten in the Clinical Bacteriology Department (Amsterdam UMC) for conducting the antimicrobial resistance testing, Eric Johnson in SNPsaurus for SMRT Sequencing, M. P. Kwint and R. Derks (Department of Human Genetics, Radboudumc, Nijmegen, The Netherlands) for SMRT/HiFi sequencing on the PacBio Sequel II, and John Atack (Griffith University) for sharing his expertise in studying Type I RM systems. Our work was funded by the European Union Horizon2020 grant 727966 (PIGSs).

